# Dietary molybdenum cofactor promotes fitness by increasing Moco content and sulfite oxidase activity in the nematode *C. elegans*

**DOI:** 10.1101/2022.08.31.506081

**Authors:** Kevin D. Oliphant, Robin R. Fettig, Jennifer Snoozy, Ralf R. Mendel, Kurt Warnhoff

## Abstract

Molybdenum cofactor (Moco) is a prosthetic group necessary for the activity of 4 unique enzymes, including the essential sulfite oxidase (SUOX-1). Moco is required for life; humans with inactivating mutations in the genes encoding Moco-biosynthetic enzymes display Moco deficiency, a rare and lethal inborn error of metabolism. Despite its importance to human health, little is known about how Moco moves among and between cells, tissues, and organisms. The prevailing view is that cells that require Moco must synthesize Moco *de novo.* Although, the nematode *Caenorhabditis elegans* appears to be an exception to this rule and has emerged as a valuable system for understanding fundamental Moco biology. *C. elegans* has the seemingly unique capacity to both synthesize its own Moco as well as acquire Moco from its microbial diet. However, the relative contribution of Moco from the diet or endogenous synthesis has not been rigorously evaluated or quantified biochemically. We genetically removed dietary or endogenous Moco sources in *C. elegans* and biochemically determined their impact on animal Moco content and SUOX-1 activity. We demonstrate that dietary Moco deficiency dramatically reduces both animal Moco content and SUOX-1 activity. Furthermore, these biochemical deficiencies have physiological consequences; we show that dietary Moco deficiency alone causes sensitivity to sulfite, the toxic substrate of SUOX-1. This work establishes the biochemical consequences of depleting dietary Moco or endogenous Moco synthesis in *C. elegans* and quantifies the surprising contribution of the diet to maintaining Moco homeostasis in *C. elegans.*

## Introduction

Molybdenum cofactor (Moco) is a prosthetic group that was present in the last universal common ancestor and continues to be synthesized in all domains of life by a conserved biosynthetic pathway (**Fig. 1**) (1,2). Moco supports the activity of 4 animal enzymes: sulfite oxidase, xanthine dehydrogenase, aldehyde oxidase, and mitochondrial amidoxime reducing component (3,4). Thus, Moco is essential to support core metabolic pathways such as sulfur amino acid and purine metabolism.

**Figure 1:**
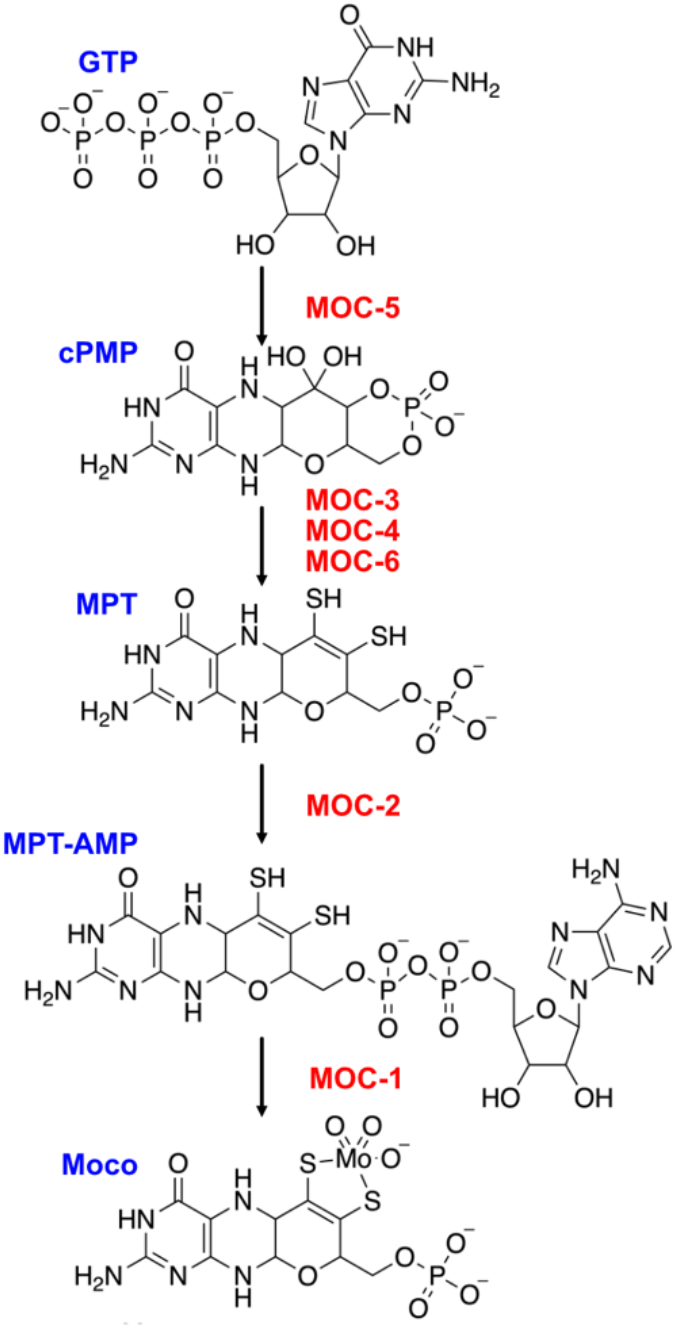
Moco biosynthesis in *C. elegans*. The *C. elegans* Moco biosynthetic pathway is displayed (enzymes, red). Structures of the following Moco-biosynthetic intermediates are displayed: guanosine triphosphate (GPT), cyclic pyranopterin monophosphate (cPMP), molybdopterin (MPT), molybdopterin adenosine monophosphate (MPT-AMP), and molybdenum cofactor (Moco).

Moco was initially revealed in the 1960s through elegant genetic studies in the fungus *Aspergillus nidulans* (5,6). Chemical mutagenesis screens were performed, leading to the isolation of mutants defective in both nitrate reductase and xanthine dehydrogenase activity. These mutants affected multiple genetic loci outlining the genes necessary for synthesizing CNX, a <u>c</u>ofactor common to <u>n</u>itrate reductase and <u>x</u>anthine dehydrogenase (6). CNX is now known as Moco.

Paralleling the genetic understanding of Moco deficiency in *A. nidulans,* human Moco deficiency was first documented in 1978 as a combined deficiency of xanthine dehydrogenase and sulfite oxidase, two Moco-requiring enzymes (7). Causative genetic lesions in human patients affected the orthologous biosynthetic pathway identified initially in *A. nidulans.* Patients presented with a severe range of symptoms as neonates, including difficulty feeding, neurological dysfunction, and eye lens dislocation. Symptoms typically appear within the first week of life, and patients generally die within three years of birth (8). Excitingly, a therapy has been developed to treat a subset of patients suffering from Moco deficiency (9). Patients with defects in cPMP synthesis can receive supplemental cPMP, which is converted to Moco by the healthy downstream Moco biosynthetic machinery. Unfortunately, cPMP therapy is only effective for patients with mutations in the first step in Moco synthesis. Supplementation with mature Moco seems a logical therapy to treat all forms of Moco deficiency. However, the instability and sensitivity of Moco to oxidation have proven experimentally limiting and precluded it from therapeutic consideration. Moco is a unique pterin and its chemical nature was ultimately resolved by studying its stable degradation products, one of which is known as “FormA” (10,11).

Given its unstable nature, very little is known about how Moco moves within and between cells. Most Moco-requiring cells are believed to synthesize Moco *de novo.* However, recent advances in the nematode *Caenorhabditis elegans* have demonstrated that Moco can move not only between cells and tissues but between disparate organisms (12–14). Unlike other animals studied, *C. elegans* with null mutations in genes encoding the Moco-biosynthetic enzymes are viable and fertile. These genes are termed *moc* in *C. elegans* for <u>mo</u>lybdenum <u>c</u>ofactor biosynthesis. *moc* mutant animals are viable because *C. elegans* stably acquires mature Moco from its bacterial diet. This dietary Moco is stabilized by Moco-binding/using proteins and is transported by unknown mechanisms throughout the *C. elegans* animal to support the activity of its Moco-requiring enzymes (13). Unsurprisingly, Moco is essential for *C. elegans* viability: animals lacking both endogenous Moco synthesis and dietary Moco arrest larval development and die. This lethality is caused by the inactivity of sulfite oxidase (SUOX-1), a Moco-requiring enzyme that oxidizes the lethal toxin sulfite to sulfate (14). Thus, *C. elegans* has two sources of Moco that function redundantly to support life under standard culture conditions: endogenous synthesis from GTP (**Fig 1**) and acquisition from its microbial diet. However, the relative importance of these distinct Moco sources has not been rigorously evaluated.

Here, we adapt biochemical protocols to quantify the contribution of dietary Moco vs. endogenously synthesized Moco. We use genetic strategies to remove either source of Moco and evaluate the impact on i) the activity of the Moco-requiring SUOX-1 enzyme and ii) *C. elegans* Moco content found in crude *C. elegans* extracts. Furthermore, we evaluate the physiological impact of removing dietary Moco vs. endogenously synthesized Moco by examining growth and development under pharmacological and genetic conditions where sulfite concentrations are toxic to animal growth. We demonstrate that dietary Moco and endogenously synthesized Moco are non-redundantly promoting Moco accumulation and SUOX-1 activity. Furthermore, removing either source of Moco causes sulfite sensitivity, indicating that both dietary Moco and endogenous Moco synthesis are required to promote optimal *C. elegans* fitness.

## Results

### Biochemical quantification of *C. elegans* SUOX-1 activity and Moco content

To biochemically evaluate *C. elegans* Moco homeostasis, we generated crude extracts from large cultures of young-adult animals. Animals were cultured on solid agar media seeded with a monoculture of *E. coli*, the food source of *C. elegans* cultivated in the laboratory. Young-adult animals were collected, extensively washed to remove *E. coli* cells, and then snap-frozen in liquid nitrogen. Samples were subsequently lysed using a bead beater, and protein concentrations were determined to allow for downstream normalization.

These crude extracts were then used to determine SUOX-1 activity and Moco content. We first analyzed samples that we anticipated would have maximal and minimal SUOX-1 activity and Moco content to validate these approaches. We reasoned that wild-type *C. elegans* fed wild-type (Moco+) *E. coli* would have high levels of SUOX-1 activity and Moco as both endogenous Moco synthesis and dietary Moco uptake are functional in this context. In contrast, we used the *moc-4; cdo-1* double mutant *C. elegans* fed *ΔmoaA* mutant (Moco-) *E. coli* as the sample where we anticipated little to no SUOX-1 activity or Moco content. *moc-4; cdo-1* double mutant *C. elegans* cannot synthesize their own Moco due to the null mutation in the *moc-4* gene that encodes molybdopterin synthase (**Fig. 1**). *moc-4* mutant animals typically display 100% penetrant larval arrest and death when cultured on Moco- *E. coli.* However, this lethality can be suppressed by mutations in *cdo-1,* which encodes cysteine dioxygenase (14). CDO-1 is a critical enzyme in the catabolism of sulfur amino acids and promotes the production of sulfites, the critical toxin when Moco is absent (14–17). *cdo-1* inactivation prevents the accumulation of sulfites and allows *C. elegans* animals to grow in the absence of endogenous and dietary Moco sources. Thus, we speculated that *moc-4; cdo-1* double mutant animals cultured on Moco- *E. coli* would be completely Moco deficient and lack SUOX-1 activity.

To determine SUOX-1 activity of these samples, we modified an assay based on a photometrically quantifiable reduction of cytochrome c (**Fig. 2A**) (18). In this reaction, sulfite is converted to sulfate, and cytochrome c acts as an electron donor. The resulting reduced form of cytochrome c has an increased absorption at OD_550,_ which we detect and quantify. Aligning with our expectations, we readily detected SUOX-1 activity from extracts of wild-type *C. elegans* fed Moco+ *E. coli* (0.33 U/mg), while we were unable to detect SUOX-1 activity from *moc-4; cdo-1* double mutant animals fed Moco- *E. coli* (**Fig. 2B,C**)*. moc-6* encodes another component of the Moco-biosynthetic machinery and is necessary for *C. elegans* Moco synthesis (**Fig. 1**) (12). Supporting our findings with *moc-4; cdo-1* animals, we were unable to detect SUOX-1 activity in crude extracts from *moc-6; cdo-1* double mutant animals fed Moco- *E. coli* (**Fig. 2C**). These results align well with prior genetic data and validate this assay to quantify SUOX-1 activity from crude *C. elegans* extracts (14).

**Figure 2:**
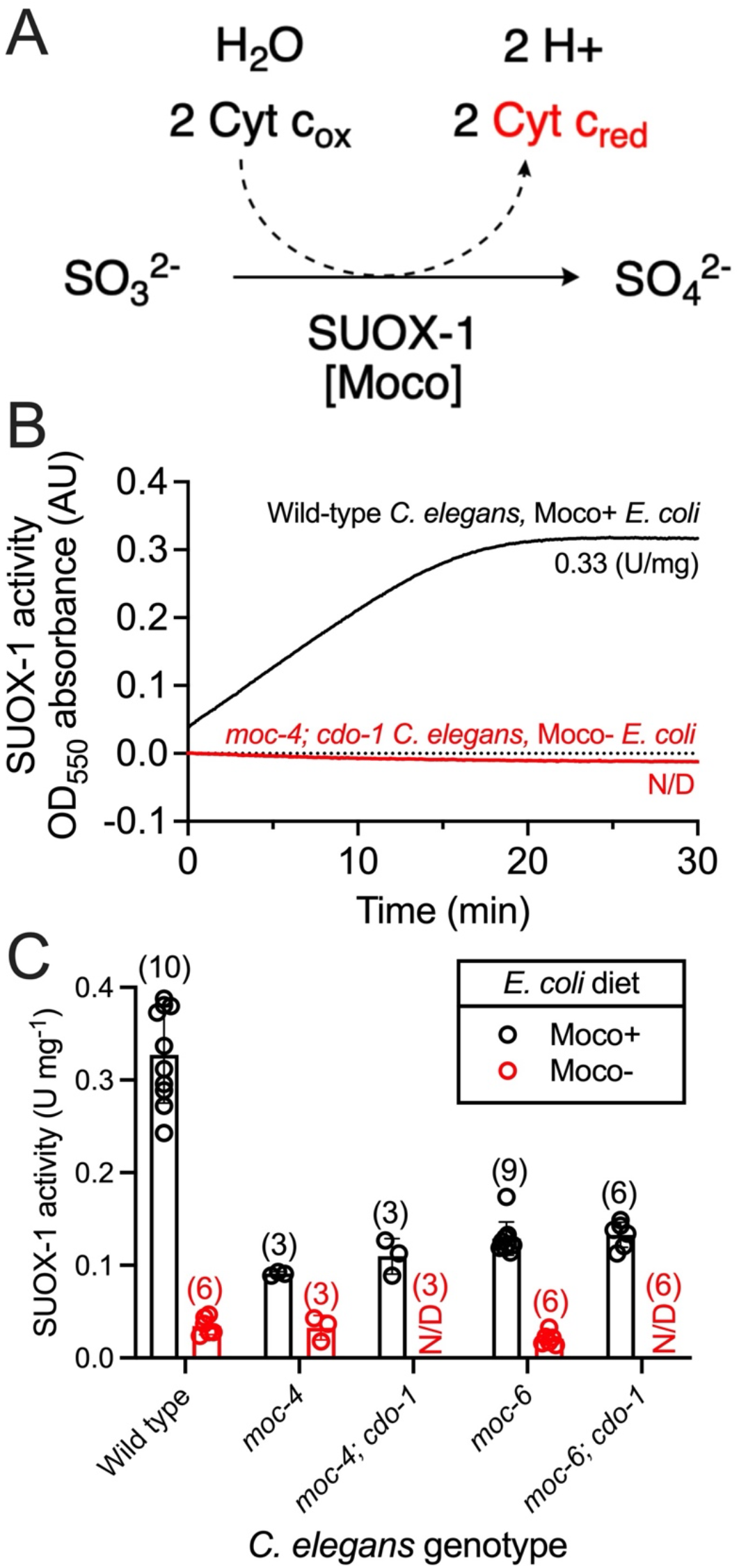
Dietary Moco and endogenous Moco synthesis promote SUOX-1 activity in *C. elegans.* A) Simplified reaction mechanism for SUOX-1 by which sulfite (SO_3_^2−^) is oxidized to sulfate (SO_4_^2−^). Sulfite-dependent SUOX-1 activity is detected via the concomitant reduction of cytochrome c. B) SUOX-1 activity was detected in crude extracts from wild-type *C. elegans* fed wild-type (Moco+) *E. coli* (black) or *moc-4(ok2571)*; *cdo-1(mg622)* mutant *C. elegans* fed *ΔmoaA* (Moco-) *E. coli* (red). Units of SUOX-1 activity per mg of protein are displayed. C) SUOX-1 activity is displayed for wild-type and mutant *C. elegans* fed either wild-type (Moco+, black) or *ΔmoaA* (Moco-, red) *E. coli.* All data points, the sample size, mean, and standard deviation are displayed for each condition. N/D indicates no SUOX-1 activity was detected in any sample.

To determine Moco content of these same samples, crude extracts were oxidized and subsequently treated with alkaline phosphatase. Iodine-dependent oxidation and dephosphorylation convert highly-unstable Moco and MPT to the stable and fluorescent derivative, dephospho-FormA (_dp_FormA) (**Fig. 3A**) (10). Oxidized metabolites were then analyzed via high-pressure liquid chromatography (HPLC) (19). _dp_FormA was detected with an average elution time of 5:40 minutes and quantified using established methods and standards (20). Supporting our genetic hypotheses, we readily detected _dp_FormA in crude extracts of wild-type *C. elegans* fed Moco+ *E. coli* (1.3 pmol/mg) and were not able to detect Moco in *moc-4; cdo-1* mutant animals fed Moco- *E. coli* (**Fig. 3B,C**). We also analyzed crude extracts from *moc-6; cdo-1* double mutant animals fed Moco- *E. coli* and were unable to detect Moco-derived _dp_FormA (**Fig. 3C**). To further validate that the peak highlighted in Fig. 3B is _dp_FormA, the same samples were prepped and analyzed without alkaline phosphatase treatment. To be detected on a reversed-phase column, FormA must be dephosphorylated, or it will not interact with the solid phase and elute in the void volume. Without alkaline phosphatase treatment, we did not see the highlighted peak in any of our samples, supporting our claim that this peak is _dp_FormA (**Fig. S1**). These results align well with expectations and validate this assay for quantifying Moco content from crude *C. elegans* extracts.

**Figure 3:**
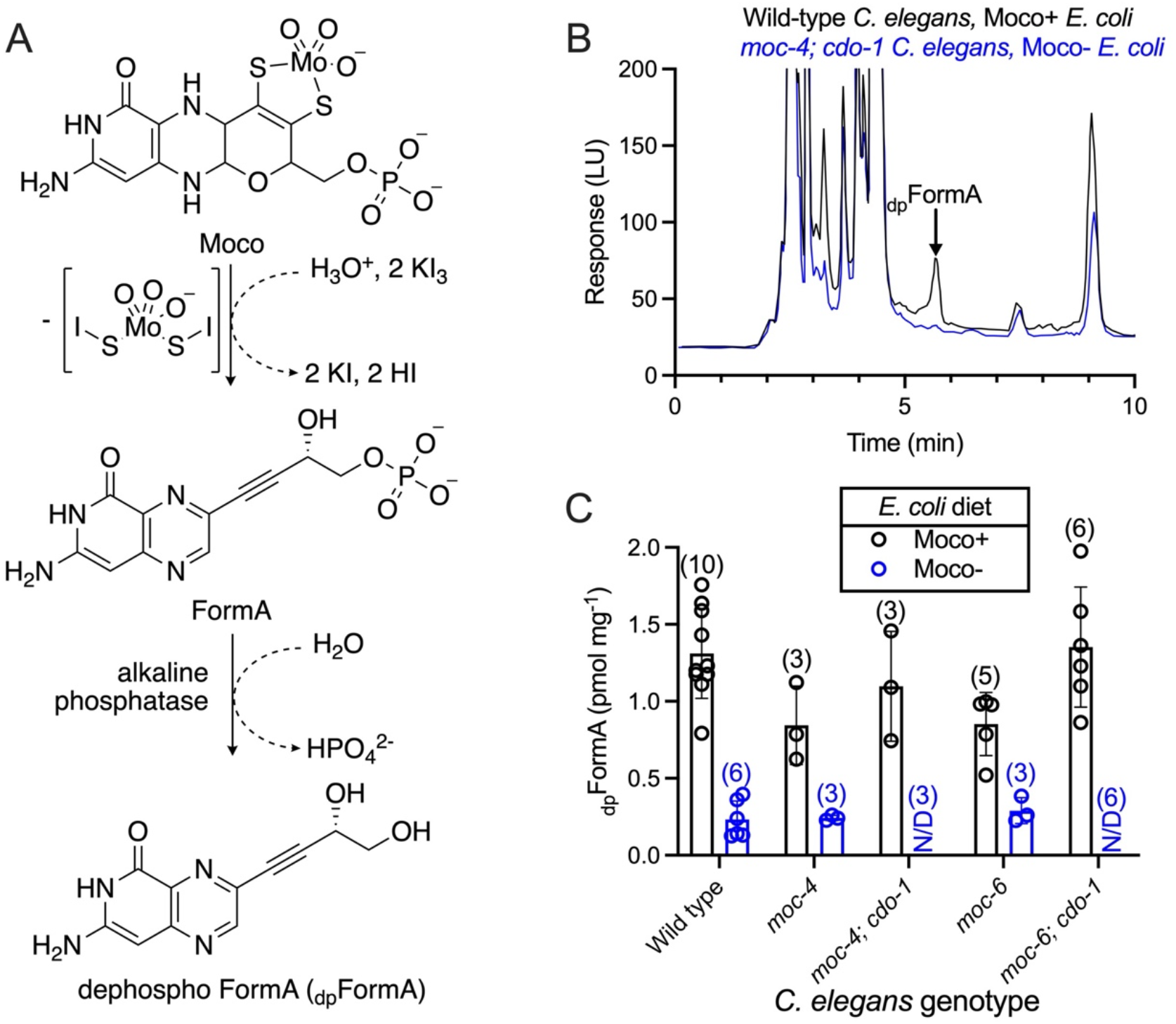
Dietary Moco and endogenous Moco synthesis promote Moco accumulation in *C. elegans.* A) A schematic of the conversion of Moco to dephospho-FormA (_dp_FormA) in the presence of an acidic iodine environment and treatment with alkaline phosphatase. B) HPLC measurements of Moco-derived _dp_FormA from crude extracts of wild-type *C. elegans* fed wild-type (Moco+) *E. coli* (black) or *moc-4(ok2571)*; *cdo-1(mg622)* mutant *C. elegans* fed *ΔmoaA* (Moco-) *E. coli* (blue). The _dp_FormA peak is indicated (black arrow). C) _dp_FormA content is displayed for wild-type, and mutant *C. elegans* fed either wild-type (Moco+, black) or *ΔmoaA* (Moco-, blue) *E. coli.* All data points, the sample size, mean, and standard deviation are displayed for each condition. N/D indicates no _dp_FormA was detected in any sample.

### Dietary Moco is necessary to promote SUOX-1 activity and Moco content in *C. elegans*

Having established new methodology for quantifying SUOX-1 activity and Moco content in *C. elegans,* we sought to address some fundamental questions regarding Moco homeostasis. Principally, how much of the total Moco content of *C. elegans* is derived from the diet as compared to endogenous synthesis? To address this question, we generated *C. elegans* samples where we deprived the animals of either i) dietary Moco (wild-type *C. elegans* fed Moco- *E. coli*) or ii) endogenous Moco synthesis (*moc-4* mutant *C. elegans* fed Moco+ *E. coli*). Extracts were generated from these samples, and SUOX-1 activity and Moco content were determined.

Wild-type *C. elegans* fed Moco- *E. coli* displayed only 11% SUOX-1 activity compared to wild-type *C. elegans* fed a Moco+ diet (**Fig. 2C**). This surprising result demonstrates that dietary Moco is necessary to promote SUOX-1 activity in wild-type *C. elegans*. Similar results were observed when we analyzed the same samples for Moco content. Wild-type *C. elegans* fed Moco- *E. coli* displayed only 18% Moco content when compared to the same animals fed a Moco+ diet (**Fig. 3C**). These data demonstrate that dietary Moco is necessary to promote Moco accumulation in wild-type *C. elegans.*

### *C. elegans* Moco biosynthesis is necessary to promote SUOX-1 activity and Moco content

Given the critical role of dietary Moco in supporting SUOX-1 activity and animal Moco content, we wondered about the relative importance of endogenous Moco synthesis. To evaluate this, we cultured *moc-4* mutant *C. elegans* on Moco+ *E. coli,* restricting the animal’s Moco source to the diet. We generated crude extracts from these samples and determined SUOX-1 activity and Moco content. *moc-4* mutant animals had 28% of the SUOX-1 activity displayed by their wild-type counterparts and 64% of the Moco content (**Fig. 2C** and **Fig. 3C**). We saw similar results when evaluating *moc-6* mutant animals fed Moco+ *E. coli* which displayed 39% of wild-type SUOX-1 activity and 65% of Moco content (**Fig. 2C** and **Fig. 3C**). These data demonstrate that endogenous Moco synthesis is necessary to promote SUOX-1 activity and Moco accumulation in *C. elegans.*

### The *C. elegans suox-1(gk738847)* mutation causes reduced SUOX-1 activity and sulfite sensitivity

SUOX-1 is an essential enzyme in both *C. elegans* and humans (14,21). To evaluate the role of *suox-1* in *C. elegans* biology, we use the hypomorphic *suox-1(gk738847)* allele as a genetic tool as it reduces *suox-1* function but does not cause overt developmental phenotypes (13,14,22). *gk738847* is a missense mutation resulting in the amino acid substitution D391N. Aspartic acid 391 is highly conserved from *C. elegans* to humans (23). To characterize the impact of the D391N substitution to SUOX-1 activity, we prepared crude extracts from young adult wild-type and *suox-1(gk738847)* animals cultured on Moco+ *E. coli.* SUOX-1 activity of these extracts was then evaluated. Our data demonstrate that *suox-1(gk738847)* mutant animals displayed 4% SUOX-1 activity compared to their wild-type counterparts (**Fig. 4A**). Despite this dramatic reduction in SUOX-1 activity, *suox-1(gk738847)* mutant animals appear superficially wild type when cultured under standard laboratory conditions. To probe for fitness defects caused by reduced SUOX-1 activity, we cultured wild-type and *suox-1(gk738847)* animals on Moco+ bacteria exposed to various concentrations of supplemental sulfite. Wild-type *C. elegans* fed Moco+ bacteria tolerate high sulfite well, displaying a half-maximal inhibitory concentration (IC_50_) of 0.0077M supplemental sulfite. In contrast, *suox-1(gk738847)* mutant animals are sensitive to sulfite displaying an IC_50_ of 0.0024M supplemental sulfite (**Fig 4B,D**). These results align well with previous studies (14). Taken together, these data biochemically demonstrate the effect of the *suox-1(gk738847)* D391N mutation on the activity of SUOX-1 in crude *C. elegans* extracts. Furthermore, we show that *suox-1* is essential for tolerating high supplemental sulfite and establish a sulfite-sensitivity assay for detecting physiological outcomes of reduced SUOX-1 activity.

**Figure 4:**
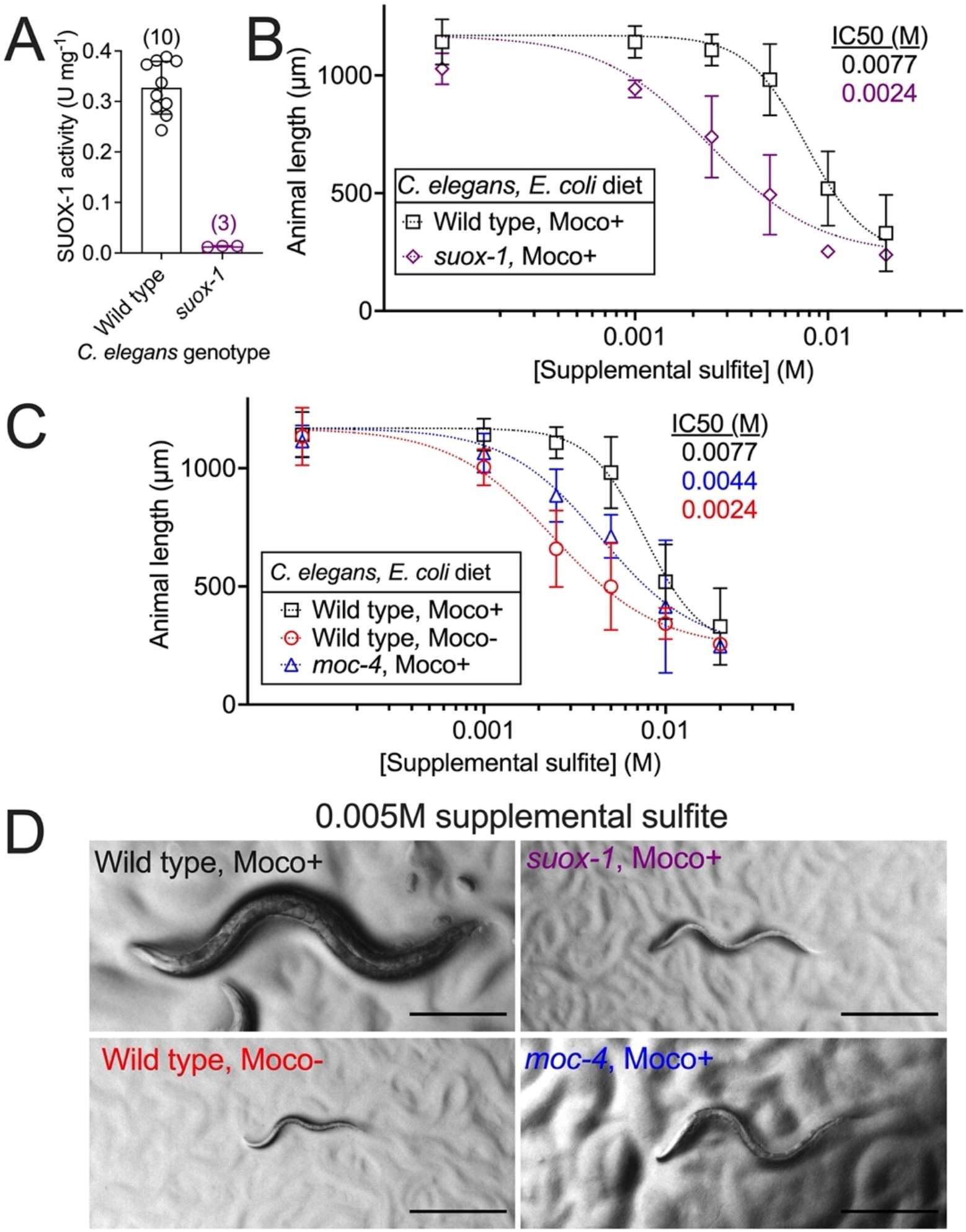
Dietary Moco and endogenous Moco synthesis are non-redundantly required for sulfite tolerance. A) SUOX-1 activity is displayed for wild-type and *suox-1(gk738847)* mutant *C. elegans* fed wild-type (Moco+) *E. coli.* All data points, the sample size, mean, and standard deviation are displayed for each condition. Note, the data displayed for wild-type *C. elegans* fed Moco+ *E. coli* are identical to the same control in Fig. 2C. B,C) Wild-type and mutant *C. elegans* were synchronized at the first stage of larval development (L1) and cultured on wild-type (Moco+) or *ΔmoaA* mutant (Moco-) *E. coli* supplemented with various concentrations of sulfite (0, 0.0001, 0.001, 0.0025, 0.005, 0.01, and 0.02M). Animal length was measured after 72 hours of growth at 20°C. Data points represent the average of 3 (*suox-1,* Moco+, and *moc-4,* Moco+) or 4 (Wild-type, Moco+, and Wild type, Moco-) biological replicates. For each biological replicate, 15 or more individual *C. elegans* animals were imaged and measured at each sulfite concentration. Error bars display standard deviation. IC_50_ values are displayed: wild-type *C. elegans* fed Moco+ *E. coli* (black), *suox-1* mutant *C. elegans* fed Moco+ *E. coli* (purple), wild-type *C. elegans* fed Moco- *E. coli* (red), and *moc-4* mutant *C. elegans* fed Moco+ *E. coli* (blue). Note that the data displayed for wild-type *C. elegans* fed Moco+ *E. coli* in Fig. 4C are identical to the same control displayed in Fig. 4B. D) Representative individuals are displayed at the critical 0.005M supplemental sulfite concentration. Scale bar is 250μm.

### Dietary Moco and endogenous Moco synthesis are non-redundantly required for sulfite tolerance

Our biochemical analyses demonstrate that wild-type *C. elegans* fed a diet of Moco- *E. coli* show decreased SUOX-1 activity **(Fig. 2C**). Like *suox-1(gk738847)* mutant animals, wild-type *C. elegans* provided Moco- *E. coli* appear superficially healthy under standard culture conditions (14). We hypothesized that the reductions in SUOX-1 activity caused by a Moco- diet would cause sulfite sensitivity. To test this, we exposed wild-type *C. elegans* cultured on Moco+ or Moco- *E. coli* to various concentrations of supplemental sulfite and analyzed growth and development. Wild-type *C. elegans* fed Moco+ *E. coli* tolerate sulfite well and display an IC_50_ of 0.0077M supplemental sulfite. By contrast, wild-type animals fed Moco- *E. coli* were sensitive to supplemental sulfite, displaying an IC_50_ of 0.0024M supplemental sulfite (**Fig. 4C,D**). These data demonstrate that dietary Moco is necessary for sulfite tolerance in *C. elegans.*

Given the requirement of dietary Moco to support sulfite tolerance, we wondered if endogenous Moco synthesis would also be essential for this process. To test the role of endogenously synthesized Moco in sulfite tolerance, we analyzed *moc-4* null mutant *C. elegans* that cannot synthesize Moco (14). We exposed *moc-4* mutant *C. elegans* cultured on Moco+ *E. coli* to various concentrations of supplemental sulfite and analyzed their growth and development. When compared to wild-type animals fed Moco+ bacteria, we found that *moc-4* mutant *C. elegans* grown on Moco+ *E. coli* were sensitive to supplemental sulfite, displaying an IC_50_ of 0.0044M supplemental sulfite (**Fig. 4C,D**). Taken together, these data demonstrate that both dietary Moco and endogenous Moco synthesis are non-redundantly required for *C. elegans* to tolerate high environmental sulfite.

### Dietary Moco is essential for the development of *suox-1(gk738847)* mutant *C. elegans*

Our biochemical analyses of Moco content and SUOX-1 activity paired with our pharmacological experiments with supplemental sulfite suggest a model for Moco homeostasis whereby both endogenous synthesis and dietary acquisition of Moco are acting non-redundantly to promote healthy Moco content in *C. elegans,* thus promoting SUOX-1 activity. To further test this model, we used the *suox-1(gk738847)* allele as a sensitized genetic background. We observed the development of these mutant animals when we altered i) dietary Moco availability and ii) endogenous Moco synthesis.

To test the role dietary Moco plays in supporting the development of *suox-1(gk738847)* mutant *C. elegans*, we cultured *suox-1(gk738847)* mutant animals on Moco+, Moco-, and mixtures of Moco+/Moco- *E. coli. suox-1(gk738847)* mutant animals grew well on Moco+ *E. coli* and 50:50 mixtures of Moco+ and Moco- *E. coli* (**Fig. 5**). However, we began to see developmental defects in *suox-1(gk738847)* mutant animals when the fraction of Moco+ *E. coli* dropped below 0.25 (**Fig. S2**). Furthermore, *suox-1(gk738847)* animals displayed the most significant reduction in growth and development when cultured on completely Moco- *E. coli* (**Fig. 5, Fig. S2**). Importantly, wild-type *C. elegans* grew well on all mixtures of Moco+ and Moco- *E. coli* (**Fig. S2**). These data align well with our previous studies and demonstrate that dietary Moco is required for the growth and development of *suox-1(gk738847)* mutant animals (14).

**Figure 5:**
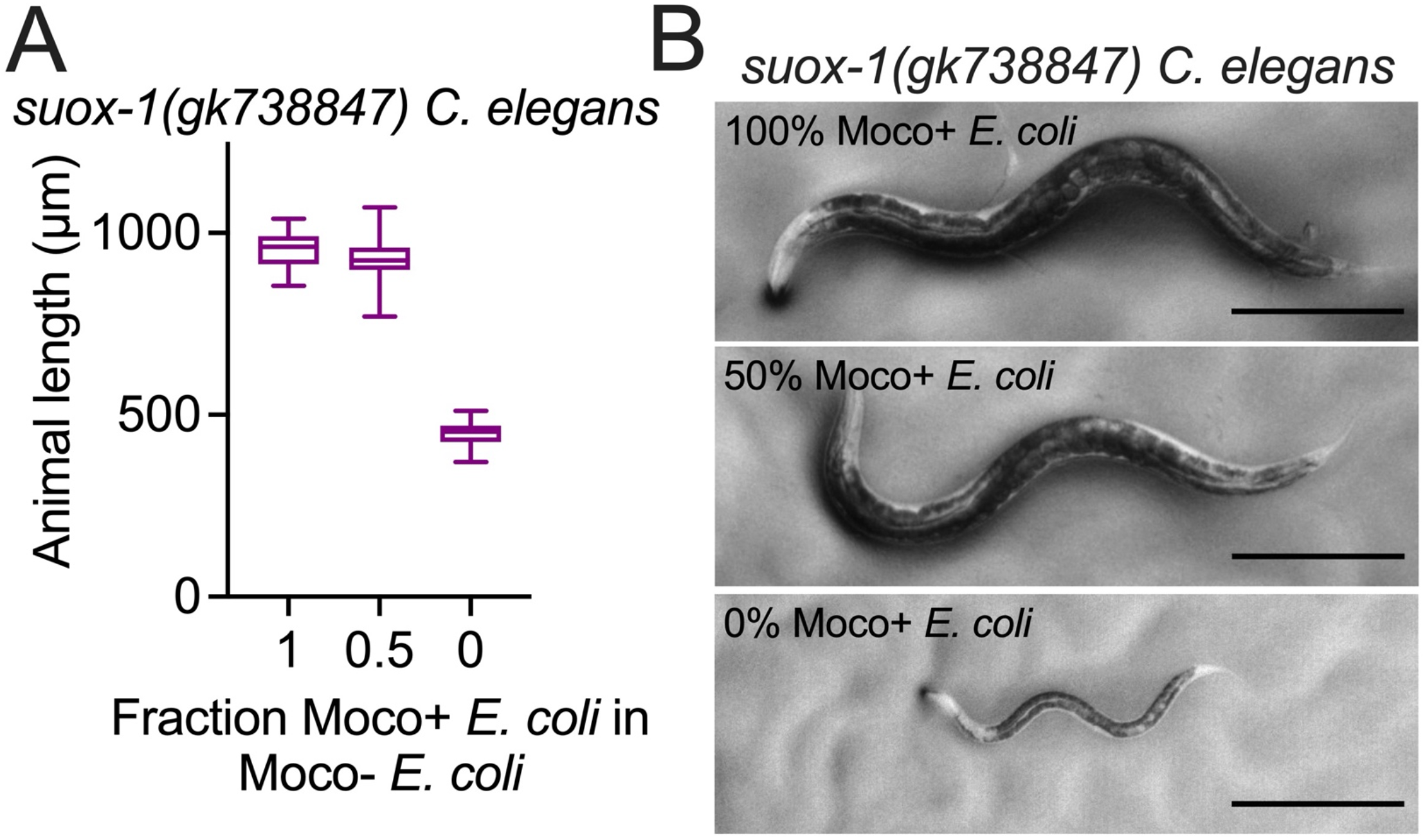
Dietary Moco is essential for *C. elegans* development when *suox-1* activity is compromised. A) *suox-1(gk738847)* mutant *C. elegans* were synchronized at the L1 stage and cultured on mixtures of wild-type (Moco+) and *ΔmoaA* mutant (Moco-) *E. coli*. Animal length was measured after 72 hours of growth at 20°C. Box plots display the median, upper, and lower quartiles, while whiskers indicate minimum and maximum data points. Sample size is 15 individuals per experiment. B) Representative individuals from Fig. 5A are displayed. Scale bar is 250μm.

### Endogenous Moco synthesis is essential for the development of *suox-1(gk738847)* mutant *C. elegans*

We then sought to test the role of endogenous Moco synthesis in supporting the growth and development of *suox-1(gk738847)* mutant *C. elegans.* To test this, we attempted to engineer double mutant strains of *C. elegans* by combining the *suox-1(gk738847)* allele with various mutations in *C. elegans* Moco-biosynthetic enzymes (*moc-1(ok366), moc-4(ok2571), moc-5(mg589),* and *moc-6(rae296)*) (12,14). In our attempts to construct these double mutants, we were unable to isolate viable double mutant strains. Furthermore, dead larvae were observed in all stages of mutant construction where we would have expected the double mutant to emerge. Thus, we hypothesized a synthetic lethal interaction between mutations in *moc* genes and the *suox-1(gk738847)* hypomorphic allele. To test this hypothesis, we constructed a strain where *suox-1(gk738847)* was homozygous, and the *moc-4(ok2571)* allele was balanced by the *tmC18* balancer (24). Given this strain, we could synchronize animals at the L1 stage and unequivocally evaluate the growth and development of *moc-4(-); suox-1(gk738847),* and *moc-4(+)/moc-4(-); suox-1(gk738847)* animals and compare their growth to *suox-1(gk738847)* single mutant animals. These mutant *C. elegans* were fed Moco+ *E. coli* throughout the experiment. Consistent with our earlier failed efforts to generate *moc; suox-1(gk738847)* double mutant animals, *moc-4(-); suox-1(gk738847)* double mutant *C. elegans* derived from the balancer strain grew extremely slowly and displayed larval lethality. By contrast, *moc-4(+)/moc-4(-); suox-1(gk738847)* and *suox-1(gk738847)* mutant animals grew well (**Fig. 6**). These data demonstrate that when *suox-1* activity is reduced via the *gk738847* mutation, endogenous Moco synthesis is essential. These results support a model whereby dietary Moco and endogenous Moco synthesis are non-redundantly promoting Moco accumulation in *C. elegans.* Increased Moco content is likely directly promoting SUOX-1 activity, which promotes sulfite tolerance (**Fig. 7**).

**Figure 6:**
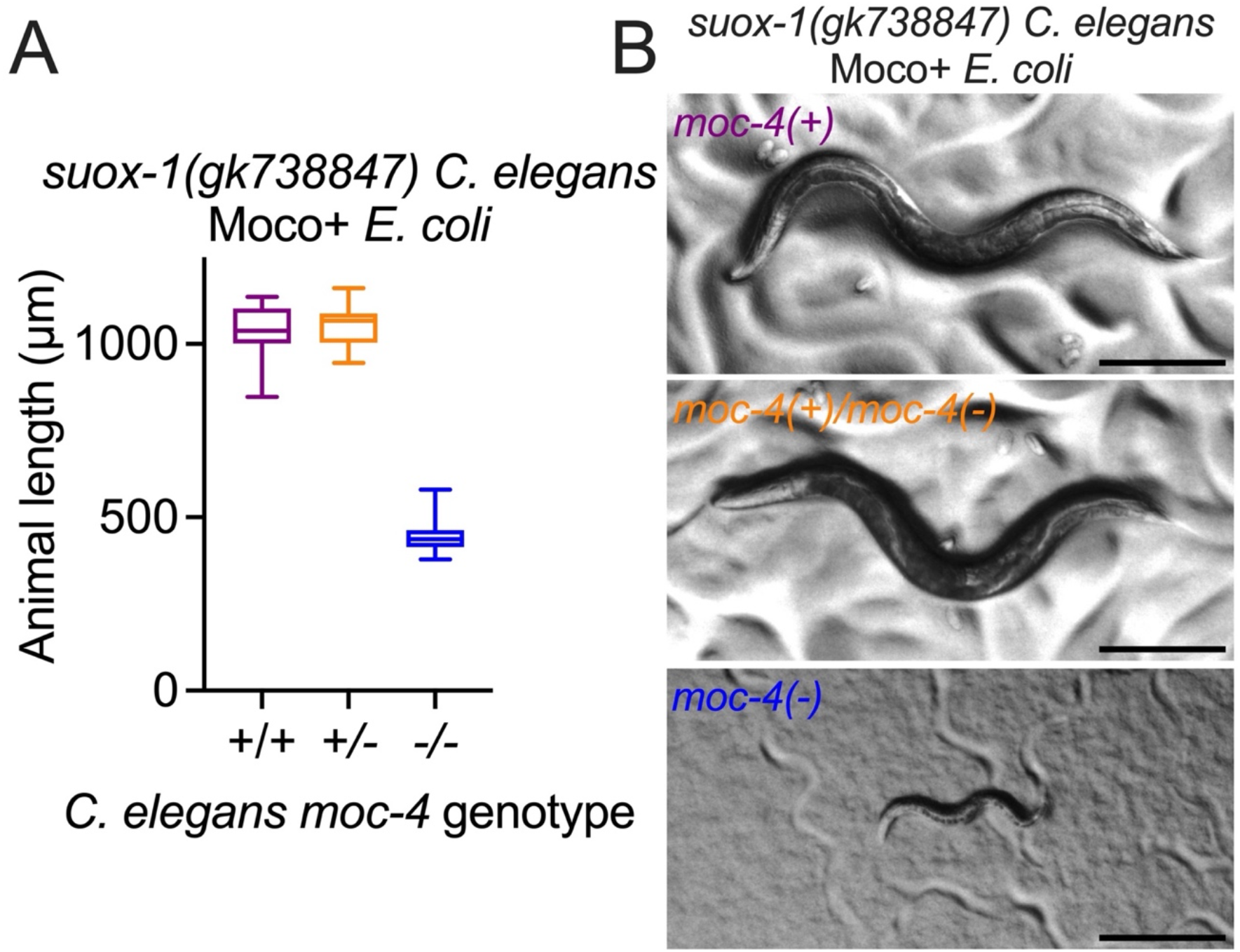
*C. elegans* Moco synthesis is essential when *suox-1* activity is compromised. A) *suox-1(gk738847), moc-4(+)/moc-4(ok2571); suox-1(gk738847),* and *moc-4(ok2571); suox-1(gk738847)* mutant *C. elegans* were synchronized at the L1 stage and cultured on wild-type (Moco+) *E. coli.* Animal length was measured after 72 hours of growth at 20°C. Box plots display the median, upper, and lower quartiles, while whiskers indicate minimum and maximum data points. Sample size is 15 individuals per experiment. Note, because *moc-4; suox-1* double mutant animals are not viable, they (along with *moc-4(+)/moc-4(-); suox-1* animals) were derived from the balanced strain USD1011 (see Experimental procedures) (24). B) Representative individuals from Fig. 6A are displayed. Scale bar is 250μm.

**Figure 7:**
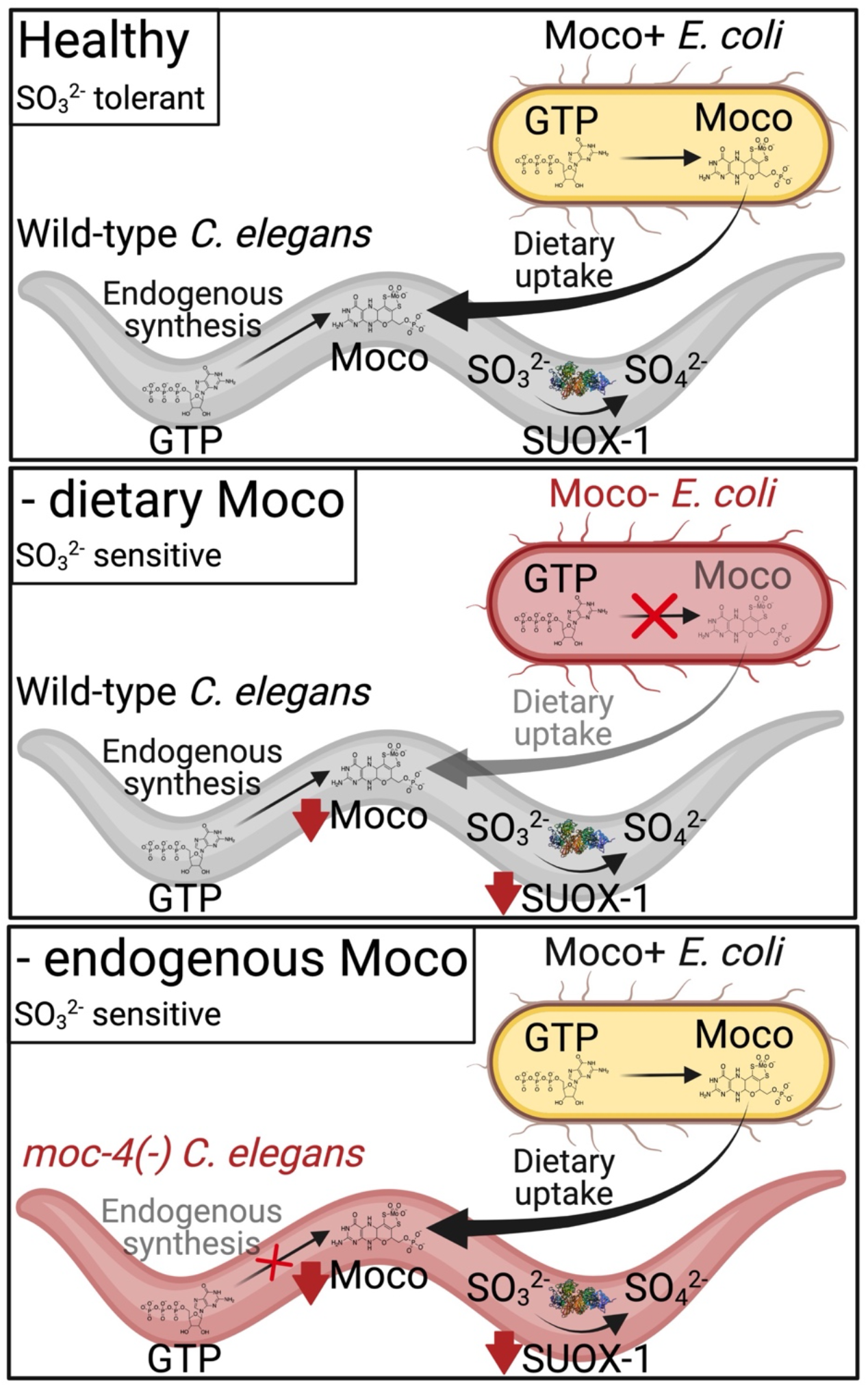
Moco homeostasis in *C. elegans*. In healthy control animals, wild-type *C. elegans* grown on wild-type *E. coli* (Moco+), SUOX-1 function and Moco content are high, leading to normal development and sulfite tolerance. SUOX-1 activity and Moco content are dramatically reduced when animals are fed a Moco- diet or when the endogenous synthesis of Moco is disrupted. These deficiencies result in sensitivity to sulfite (SO_3_^2−^).

## Discussion

### Dietary *E. coli* are a major source of Moco for *C. elegans*

The free-living nematode *C. elegans* acquires Moco from its bacterial diet or by synthesizing it *de novo* from GTP. Under standard laboratory conditions, either source of Moco is sufficient to promote growth, development, and reproduction. Thus, when growth conditions are ideal, these pathways appear to operate redundantly to support life. However, this conclusion relies exclusively on measuring the rate of development and fertility (14).

To expand our understanding of Moco homeostasis, we produced *C. elegans* extracts from animals lacking either endogenous Moco synthesis or dietary Moco. These extracts were used to quantify Moco content and SUOX-1 activity, providing biochemical clarity to our genetic system (12–14). We found that *C. elegans* mutants lacking endogenous Moco biosynthesis displayed reduced Moco content and SUOX-1 activity. *moc-4* and *moc-6* mutant *C. elegans* fed Moco+ *E. coli* showed 64% and 65% of Moco content and 28% and 39% SUOX-1 activity, respectively, compared to wild-type controls. These decreases were expected as Moco synthesis has long been understood to be an essential source of Moco for the cell.

Surprisingly, we found even more severe defects when evaluating the biochemical impacts of a Moco-deficient diet. Wild-type *C. elegans* fed a Moco- diet displayed 18% of the Moco content and 11% of SUOX-1 activity compared to wild-type animals fed a Moco+ diet. This demonstrates that the *C. elegans* diet substantially contributes to Moco homeostasis in *C. elegans* animals. Thus, dietary Moco and endogenous Moco synthesis function non-redundantly to promote Moco content and SUOX-1 activity.

Having demonstrated the biochemical impacts of either dietary or endogenous Moco loss, we wondered whether these deficiencies in Moco content or SUOX-1 activity would cause a fitness defect. To evaluate this, we employed sulfite as pharmacological intervention. Sulfite is a useful tool as it is a toxin and the primary substrate of SUOX-1. Sulfite is typically produced as a byproduct of sulfur amino acid metabolism (25). Thus, maintaining SUOX-1 activity is critical for sulfite detoxification and *C. elegans* survival. We observed that *C. elegans* lacking either source of Moco displayed sulfite sensitivity, demonstrating the non-redundant roles of dietary and endogenous Moco in promoting sulfite tolerance. Supporting this result, we found that both sources of Moco were also required for life in a *suox-1(gk738847)* hypomorphic mutant background.

Given these results, we propose the model that when food is abundant, *C. elegans* animals rely on both endogenous Moco synthesis and dietary Moco to support Moco homeostasis (**Fig. 7**). While endogenous Moco synthesis has long been established to be critical for supporting Moco homeostasis, our work demonstrates the equal importance of dietary Moco in the life and survival of *C. elegans.*

### Are two sources of Moco better than one?

Why would the *C. elegans* genome retain Moco biosynthetic machinery if the animals can acquire the cofactor so efficiently from the diet? Why is Moco not a vitamin for *C. elegans,* as is the case for other essential coenzymes (26–28)? Logic dictates that there must be a fitness advantage to maintaining both strategies of increasing cellular Moco. We speculate that acquiring Moco from dietary microbes (when they are abundant) is energetically favorable compared to *de novo* Moco synthesis from GTP. However, endogenous Moco synthesis is likely a more reliable Moco source during the life of a wild *C. elegans* individual. This is because wild *C. elegans* have a “boom-and-bust” life cycle (29). When conditions are ideal, newly hatched *C. elegans* pass through four larval stages and reach fertile adulthood in about three days. However, under stressful conditions (low food, high population density, high temperature), *C. elegans* enter an alternative non-feeding diapause state known as dauer (30). Dauer larvae are stress-resistant and can survive months as they disperse and seek new food. Wild *C. elegans* live the majority of their lives as non-feeding dauer larvae that are likely reliant on endogenous Moco synthesis (31). As dauer larvae locate food (usually microbes found on rotting fruit or plant stems) and return to favorable conditions, they resume development and reach fertile adulthood (30). During this “boom” time, dietary Moco would be abundant as ~70% of bacterial genomes encode Moco biosynthetic enzymes (2). Yet, the food source will eventually be consumed, dauer larvae will emerge in the population, and the cycle will repeat. Considering this natural history, redundant sources of Moco seem logical. When no food is available for weeks/months, the animals must rely on endogenous Moco synthesis to survive. However, once a microbial food source is identified, *C. elegans* additionally harvest Moco from their diet, promoting Moco accumulation and enzymatic function within the animals.

### Moco bioavailability in *C. elegans* and beyond

Moco is an ancient and essential prosthetic group; Moco synthesizing and requiring enzymes were present in LUCA and persist in all domains of life today (1,2). Thus, it seems likely that Moco biology uncovered in the nematode *C. elegans* will be found throughout the tree of life. Our discovery that dietary Moco promotes Moco content, SUOX-1 activity, and rescues *C. elegans* Moco deficiency lays additional intellectual groundwork for developing Moco supplementation as a therapy to treat human Moco deficiency (13,14). However, uptake of exogenous Moco has yet to be examined or observed in another organism. It is crucial to determine if human cells are competent to acquire exogenous Moco as this is a critical next step in developing therapeutic Moco.

In addition to therapeutic considerations, insights into Moco biology from the nematode *C. elegans* reveal many new fundamental questions about Moco biology: how is Moco moving from bacteria to *C. elegans*? How is Moco stably harvested from the exogenous donor proteins? Once harvested, how is Moco traversing cell membranes? Are there animal Moco chaperones facilitating these processes? Given the ancient and conserved nature of Moco, we expect pathways and phenomena uncovered in *C. elegans* to be at play across the diversity of life.

## Experimental procedures

### Animal cultivation

*C. elegans* strains were cultured using established protocols (32). Briefly, animals were cultured at 20°C on nematode growth media (NGM) seeded with wild-type *E. coli* (OP50) unless otherwise noted. The wild-type *C. elegans* strain was Bristol N2.

Additional *C. elegans* strains used in this work were:

GR2253 [*moc-4(ok2571) I*],
USD989 [*moc-4(ok2571) I; cdo-1(mg622) X*],
USD1011 [*moc-4(ok2571)/tmC18 [tmIs1236] I; suox-1(gk738847) X*],
USD955 [*moc-6(rae296) III*],
USD959 [*moc-6(rae296) III; cdo-1(mg622) X*],
GR2256 [*moc-5(mg589) X*]
GR2254 [*moc-1(ok366) X*], and
GR2269 [*suox-1(gk738847) X*].

Additional *E. coli* strains used in this work were:

BW25113 (wild type, Moco+) and
JW0764-2 (*ΔmoaA753::kan,* Moco-).

### Protein and metabolite extraction from *C. elegans*

Synchronized populations of 10,000 to 20,000 *C. elegans* animals were cultured at 20°C and harvested at the young adult stage of development. To remove dietary bacterial cells from the sample, animals were washed in excess M9 buffer three times and then incubated in M9 buffer for 1 hour at 20°C with gentle rocking. Animals were subsequently washed one additional time with excess M9 buffer, pelleted, and flash frozen in liquid nitrogen. All samples were cultured on either BW25113 (wild type, Moco+) or JW0764-2 (*ΔmoaA753::kan,* Moco-) *E. coli* for the entire experiment. However, if *moc-*mutant *C. elegans* are cultured on *ΔmoaA* (Moco-) *E. coli* from early larval stages, they undergo larval arrest and death (12,14). To allow for biochemical analyses of *moc-4* and *moc-6* single mutant animals cultured on *ΔmoaA* (Moco-) *E. coli*, these mutant strains were initially cultured on Moco+ *E. coli* until the L4 stage of development (48 hours post L1 synchronization). These L4 larvae were then collected, extensively washed as described above, and re-introduced onto new culture dishes containing NGM seeded with Moco- *E. coli.* Thus, dietary Moco was depleted at a later stage in development, as previously described (12,14).

For total protein and metabolite extraction, flash-frozen *C. elegans* samples were lysed in 400 μl of lysis buffer (20 mM HEPES, 150 mM NaCl, 1 mM EDTA, 0.5% Triton X-100, pH 7.5) using a FastPrep-24 (M.P. Biomedicals Irvine, USA) four times for 30 seconds at 6.5 m/s with 5-minute breaks and then incubated for 30 minutes on ice. Protein concentrations were determined using a Bradford protein assay (Carl Roth Karlsruhe, Germany). Absorption was measured using a Multiskan GO microplate spectrophotometer (ThermoFisher Scientific Waltham, USA).

### Detection of sulfite oxidase activity

Sulfite oxidase activity measurements were based on a sulfite-dependent enzymatic reduction of cytochrome c (18). 10 – 50 μg protein of total crude extract from *C. elegans* were used for quantification. 50 μl of each dilution were loaded into a 96-well plate. 180 μl of the SUOX-1 activity buffer (100 mM Tris-HCl, 0.1 mM EDTA, 0.04 mM cytochrome c_ox_, pH 8.5) were added to the well. Subsequently, samples were incubated for 5 minutes on ice. The reaction was initiated by adding 20 μl of a 5 mM sodium sulfite solution. As a control, 20 μl _dd_H_2_O was added instead. Immediately after adding sulfite, an absorption change at 550 nm was measured in a Multiskan GO microplate spectrophotometer (ThermoFisher Scientific Waltham, USA). The detection was set over a time course of 60 min with 720 measurements. For visualization, the _dd_H_2_O control was subtracted from the sample with sulfite, which was used to plot a time-dependent OD change. The SUOX-1 activity was determined from the slope of the linear range of the curve. 1U is defined as the amount of protein required to reach an absorption increase of 1.0 per minute.

### Quantification of Moco/MPT using FormA

Quantifying the Moco and molybdopterin (MPT) content of the crude extracts was performed by oxidizing Moco/MPT in a well-defined environment to FormA, a stable and fluorescent Moco/MPT oxidation product (10). For oxidation of Moco, 500 μg of crude protein extract in 400 μl of 100 mM Tris-HCL pH 7.2 were used. Samples were oxidized with 50 μl oxidation solution (1% I_2_, 2% KI in 1 M HCl) for 16 hours in the dark at room temperature. After oxidation, the samples were centrifuged at 10,000 x g for 10 min, and the supernatant was transferred to a new reaction tube. 55 μl 1% ascorbic acid solution were added to stop the oxidation reaction. Subsequently, 200μl 1 M Tris, 13 μl MgCl_2,_ and 2 μl alkaline phosphatase (Roche Basel, Switzerland) were added and incubated overnight in the dark at room temperature. The HPLC detection of _dp_FormA was performed at room temperature using a reversed-phase C-18 column (250 mm × 4.6 mm, 5 μm, ReproSil-Pur Basic C-18 HD) with an Agilent 1100 HPLC system containing a fluorescence detector (19). The specific parameters were set with a flow rate of 1 ml min^−1^ using an isocratic run with 5 mM ammonium acetate and 15% (v/v) methanol as the mobile phase. For the fluorescence detection λ ex = 302 nm, λ em = 451 nm were set. For data analysis, OpenLab CDS Version 2.2.0.600 was utilized. Calibration was carried out using synthetic _dp_FormA (20).

### *C. elegans* growth assays

Growth assays were performed as previously described with minor modifications (13,14). Briefly, wild-type and mutant *C. elegans* were synchronized at the first stage of larval development (L1) and subsequently cultured on NGM seeded with wild-type or mutant *E. coli* for 72 hours at 20°C. Live animals were then imaged using an SMZ25 stereomicroscope (Nikon) equipped with an ORCA-Flash4.0 camera (Hamamatsu). Images were captured using NIS-Elements software (Nikon) and processed using ImageJ. Animal length was measured from the tip of the head to the end of the tail. GraphPad Prism software was used for calculations of median, upper, and lower quartiles.

For sulfite sensitivity experiments, the dietary *E. coli* were pelleted and re-suspended with various concentrations of sodium sulfite (Millipore Sigma) in water. The *E. coli* (5X concentrated)*-*sulfite slurries were then seeded onto the NGM media as a food source for *C. elegans.* The concentration of sulfite displayed assumes the sulfite diffuses evenly throughout the 10ml of NGM media. A limitation to this method is that sulfite oxidizes to sulfate in water, and thus the presented concentrations likely overestimate the sulfite concentration experienced by *C. elegans* in these assays (33,34). To mitigate sulfite oxidation, we worked quickly to limit the time between establishment of the culture dishes (NGM, *E. coli,* sulfite) and the addition of the appropriate *C. elegans* animals. Growth assays were then performed as described above. IC_50_ measurements were calculated by nonlinear regression analysis using GraphPad Prism software.

For experiments where Moco+ and Moco- bacterial diets were mixed, BW25113 and JW0764-2 *E. coli* were first cultured overnight in LB at 37°C in a shaking incubator. Overnight cultures were then concentrated ten times. These 10X concentrated stocks were then mixed yielding various fractions of Moco+ (wild type) *E. coli* in Moco- (*ΔmoaA*) mutant *E. coli*. These *E. coli* mixtures were then seeded onto NGM supplemented with the antibiotic Streptomycin, preventing additional *E. coli* growth. Growth assays were performed as described above.

## Data availability

All data are presented within the manuscript.

## Supporting information

This article contains supporting information.

## Acknowledgments

We thank the *Caenorhabditis* Genetics Center for providing *C. elegans* strains. We thank the National Institute of Genetics, National BioResource Project (NIG, Japan) for providing *E. coli* strains.

## Supporting Information

**Supporting Figure 1:**
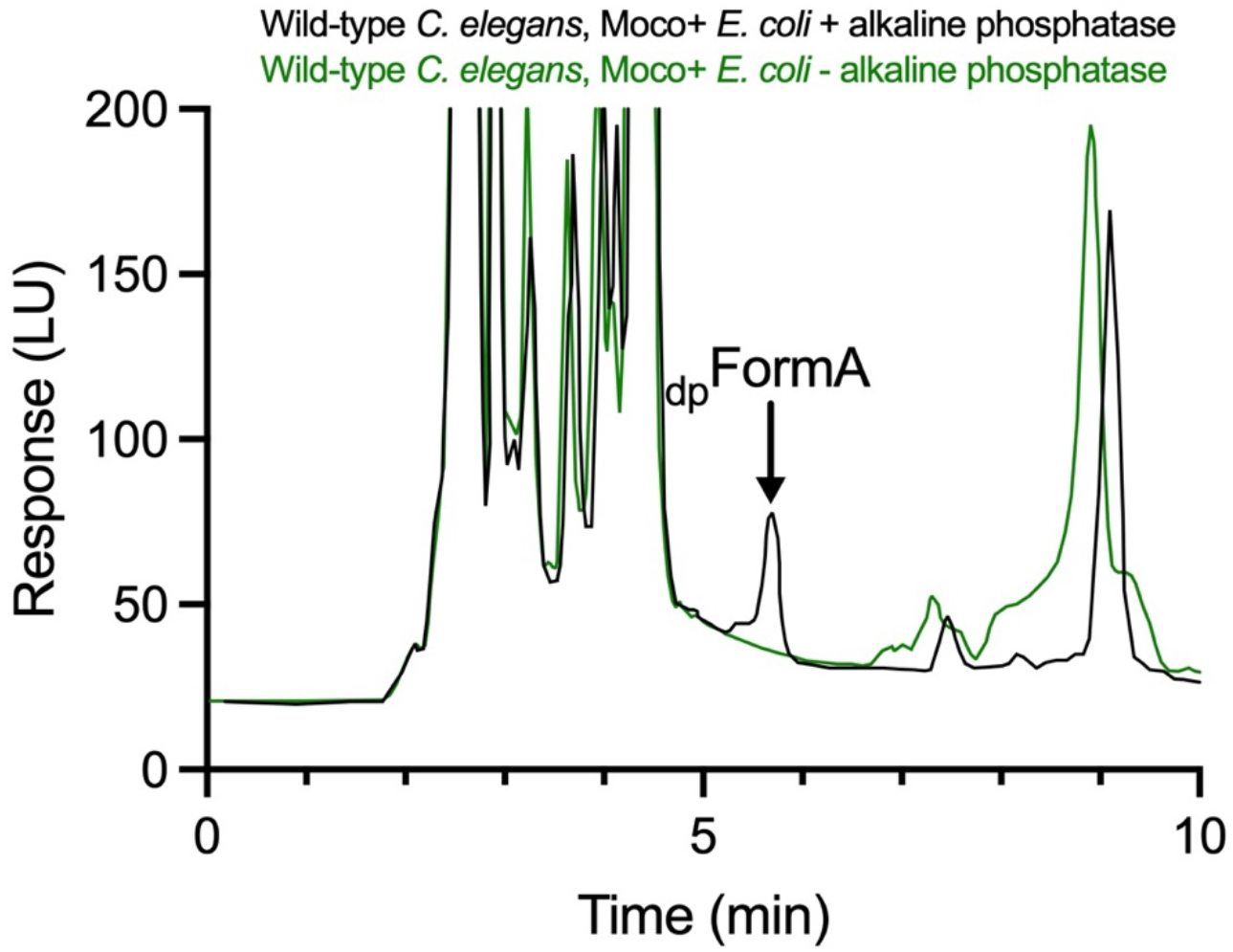
FormA analysis with and without the addition of alkaline phosphatase. HPLC measurements of Moco-derived dephospho-FormA (_dp_FormA) from crude extracts of wild-type *C. elegans* fed wild-type (Moco+) *E. coli*. Oxidized extracts were evaluated with (black) or without (green) alkaline phosphatase treatment. The _dp_FormA peak is indicated (black arrow).

**Supporting Figure 2:**
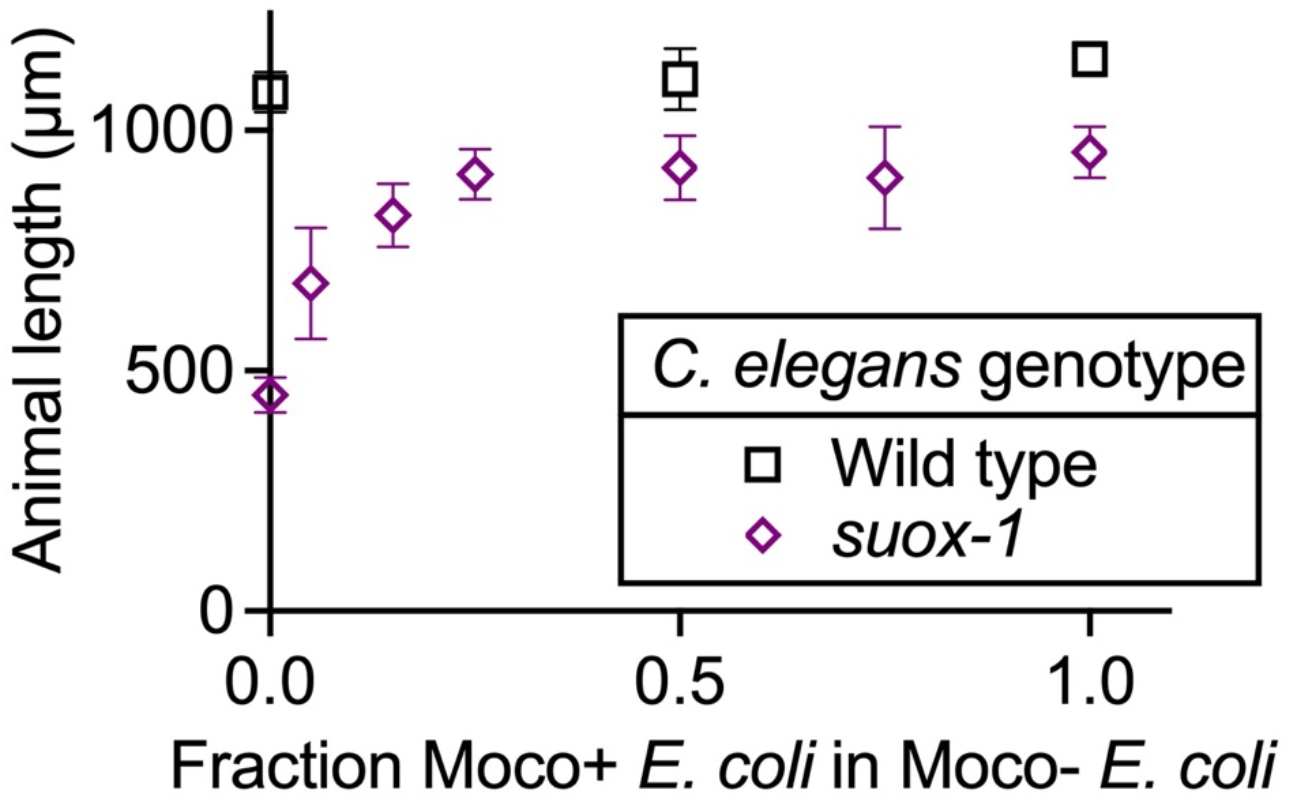
*suox-1(gk738847)* mutant *C. elegans* are sensitive to dietary Moco deficiency. Wild-type and *suox-1(gk738847)* mutant *C. elegans* were synchronized at the L1 stage and cultured on diets with different fractions of wild-type (Moco+) in *ΔmoaA* mutant (Moco-) *E. coli* (0, 0.05, 0.15, 0.25, 0.5, 0.75, 1). Animal length was measured after 72 hours of growth at 20°C. Sample size is 15 individuals per data point. Mean and standard deviation are displayed. Note, datapoints at 0, 0.5, and 1.0 fraction Moco+ in Moco- *E. coli* for *suox-1* mutant *C. elegans* are also displayed in Fig. 5A.

